# Matrix stiffness and stress relaxation regulate matrix-bound nanovesicle release from alginate hydrogels

**DOI:** 10.64898/2026.05.28.728372

**Authors:** Renata Dos Reis Marques, Jane A. Baude, Gianna M. Gathman, Ava I. Salami, Ryan S. Stowers, Marley J. Dewey

## Abstract

Matrix-bound nanovesicles (MBVs) are a recently discovered subclass of small extracellular vesicles (EVs) that reside within the extracellular matrix of non-mineralized tissues throughout the body. Functionally, MBVs exhibit unique immunomodulatory properties that have been leveraged therapeutically to treat various tissue pathologies, including periprosthetic osteolysis, rheumatoid arthritis, and skeletal muscle injury. However, like other EVs, the therapeutic efficacy of MBV applications is limited by delivery methods, namely bolus injections, that offer poor control of EV persistence and bioavailability at the site of administration. We hypothesized that a superior MBV delivery platform could be developed by entrapping MBVs in a tunable, engineered alginate matrix to control retention and release of MBVs on therapeutically relevant timescales. To this end, we encapsulated dermal fibroblast MBVs in bioinert alginate hydrogels of varying stiffness and stress relaxation rates to determine the impact of matrix mechanical properties on MBV release and retention over a 14-day period. We found that stiffer matrices increased MBV release compared to their softer counterparts. Additionally, fast-relaxing matrices exhibited release of MBVs in the first four days of release experiments, in contrast with slow-relaxing matrices, which promoted long-term sequestration of nearly all encapsulated MBVs regardless of differences in matrix stiffness. Our results offer promise that alginate hydrogels can be utilized for more precise control of MBV delivery in the body and may overcome limitations associated with current EV administration methods.

## 1. INTRODUCTION

Extracellular vesicles (EVs) are cell-secreted lipid bilayer-bound nanoparticles that encapsulate bioactive molecules such as RNAs, proteins, and lipids. EVs have been shown to mediate biological processes such as tissue homeostasis, aging, cancer progression, wound healing, and inflammation, among many others, typically through paracrine signaling. Several subpopulations of EVs have been isolated from biological fluids, including exosomes, microvesicles, and apoptotic bodies, though only recently a population of small EVs was identified within the extracellular matrix of non-mineralized tissues and *in vitro* cultures^1–3^, termed “matrix-bound nanovesicles” (MBVs). These matrix-associated EVs are tissue-specific^4^, reflect the diseased state of tissues^3^, and are characteristically distinct from biofluid-derived EVs (hereafter referred to as “liquid-EVs”). Liquid-EVs and MBVs are differentiated by their microRNA, lipidomic^5^, and proteomic^6,7^ compositions, and exert different therapeutic functions^8,9^. Moreover, MBVs exhibit immunomodulatory properties without cytotoxic effects^10^ with demonstrated efficacy in the treatment of various tissue disorders, particularly optic nerve injury^11^, periprosthetic osteolysis^12^, and pristane-induced rheumatoid arthritis^13^. Despite their promise as therapeutics, however, the efficacy of MBVs, much like other types of EVs, is compromised by rapid systemic clearance following administration that limits their bioavailability^10^.

Studies investigating EVs as therapeutics typically administer them as a bolus injection. However, the therapeutic potential of small EVs, including MBVs, is hindered by their rapid dispersal and lack of targeted delivery at the injection site due to their size (50-200 nm), resulting in premature systemic clearance from the body. Specifically, a systematic review comparing the biodistribution of EVs across multiple *in vivo* experiments noted that regardless of type, liquid-EVs localize to the liver, lungs, kidneys, and spleen and are often cleared in 1-24 hours^14^. The treatment of chronic conditions or injuries with prolonged healing timelines typically requires therapeutic signals to be sustained for much longer; for example, large-scale bone defects can require over 6 months to heal^15^, necessitating an impractical number of injections of EVs to treat these wounds. These challenges underscore the need for greater control of EV delivery and retention at the wound site to maximize therapeutic efficacy and reduce off-target effects.

Three-dimensional hydrogels offer a compelling platform for EV delivery as they have the potential to entrap EVs within a matrix, reduce systemic EV clearance, and enhance EV bioavailability. Previous studies have shown that EV interactions with materials are dependent on material mechanical properties^16–18^, material degradation^19–22^, electrostatic interactions^23,24^, and EV binding to extracellular matrix proteins^25–27^, to name a few, demonstrating that material design strategies can be used to modulate EV delivery for therapeutic applications. However, many studies incorporating EVs in hydrogels are still limited by their short-term observation periods and indirect methods for quantifying particle release (e.g. protein quantification) and often require complex functionalization strategies to promote EV attachment. Most importantly, MBVs have yet to be incorporated into biomaterials for controlled release despite their promise as immunomodulatory therapeutics. Therefore, understanding and leveraging interactions between EVs and hydrogels mediated by material properties could enable fine-tuned EV release kinetics for specific clinical applications.

Here, we utilize a mechanically tunable alginate hydrogel platform to assess the impact of material mechanical properties on MBV release and retention. Alginate is a bioinert polysaccharide extracted from brown seaweed that can form hydrogels upon ionic crosslinking. Alginate hydrogels are widely used for drug and cell delivery due to their cytocompatible gelation mechanism and their readily tunable physical properties^28–31^. Unlike synthetic hydrogels, in which mesh size and crosslink density are inversely related through defined mathematical models, alginate networks exhibit a unique structural decoupling^32^. The gelation process of alginate hydrogels can be described according to the egg-box model, as Ca^2+^ ions induce gelation by nesting into pre-existing coordination sites along the guluronate (G) blocks^33,34^. Increasing the calcium concentration primarily saturates these vacant “egg-box” sites rather than creating new junction zones. As a result, the hydrogel’s mechanical stiffness can be increased without significantly altering its mesh size, and the underlying network architecture remains relatively constant. Additionally, matrix stiffness and stress relaxation can be independently altered by adjusting crosslinker concentration and alginate molecular weight, respectively^30^; as such, this system is ideal for isolating the individual and combinatorial effects of matrix stiffness and stress relaxation on EV release kinetics. Finally, the bioinert nature of alginate minimizes confounding EV-hydrogel interactions mediated by receptor-ligand binding, further isolating the effects of material mechanical properties on MBV release. In this study, we encapsulated dermal fibroblast MBVs in alginate hydrogels of varying stiffness and stress relaxation rates and measured vesicle release and retention over a period of 14 days. We also compared these results to latex nanoparticles of similar size to MBVs to evaluate whether our release results were primarily due to size. Our study is the first to demonstrate the impact of matrix mechanical properties on MBV interactions within alginate hydrogels, establishing the feasibility of hydrogels as modular MBV delivery platforms.

## 2. MATERIALS & METHODS

### 2.1 Cell culture

Primary adult human dermal fibroblasts (41-year-old female donor, American Type Culture Collection, PCS-201-012) were acquired at passage 1. Cells were cultured in normal growth medium comprised of low glucose DMEM (Gibco, 11885084) supplemented with 10% fetal bovine serum (Genesee Scientific, 25-514) and 1% antibiotic-antimycotic (Gibco, 15240062) and maintained at 37°C and 5% CO_2_. The initial cell population was expanded until passage 4 prior to cell sheet formation.

### 2.2 Cell sheet formation

Primary human dermal fibroblasts at passage 4 were seeded at 500,000 cells per 150 mm petri dish (Genesee Scientific, 25-203) and allowed to attach overnight. The next day, normal growth medium was supplemented with 50 μM L-ascorbic acid-2-phosphate (Tokyo Chemical Industry, A2521) to induce extracellular matrix deposition. Cell sheets were cultured for a total of 21 days. To reduce possible cross-contamination of extracellular vesicles, L-ascorbic acid-containing growth medium was supplemented with extracellular vesicle-depleted fetal bovine serum 6 days prior to matrix-bound nanovesicle isolation. Extracellular vesicle-depleted fetal bovine serum was prepared by ultracentrifugation at 100,000 g for 18 hours at 4°C prior to medium exchange. The resulting supernatant was recovered and filter-sterilized through a 0.22 μm PES membrane. Extracellular vesicle-depleted fetal bovine serum was stored at −20°C until use.

### 2.3 Isolation of matrix-bound nanovesicles

After 21 days of *in vitro* cell culture, the culture medium was discarded, and cell sheets were washed with 1X PBS to remove residual medium. The cell-deposited extracellular matrix was then digested with 10 μg/mL Liberase TH enzyme (Sigma Aldrich, 5401151001) in Liberase buffer (50 mM Tris-HCl pH = 8.0, 200 mM NaCl, 5 mM CaCl_2_) for 3 hours at 37°C with constant rotation. The digested extracellular matrix was differentially centrifuged at 500 g for 10 minutes, 2,500 g for 20 minutes, and 10,000 g for 30 minutes at 4°C, then filtered through a 0.22 µm PES membrane. Extracellular matrix digests were concentrated using a 100 kDa molecular weight cut-off (MWCO) filter column (Sigma Aldrich, UFC910024) at 3,220 g (4°C) and purified with gravity size-exclusion chromatography. Disposable chromatography columns (BioRad, 7321010) were prepared using 10 mL of Sepharose CL-2B (Sigma Aldrich, CL2B300). After a thorough flush with 1X PBS, 1 mL of extracellular matrix digest was loaded into the column, and 1 mL of PBS was added to elute each fraction for a total of 10 fractions. Fractions 3 through 5 were collected, pooled, and further concentrated using 100 kDa MWCO filters at 3,220 g (4°C). Concentrated MBVs in 1X PBS were aliquoted, stored at −80°C, and were subjected to no more than two freeze-thaw cycles.

### 2.4 Quantification of size and concentration of isolated matrix-bound nanovesicles and latex nanoparticles

Shortly after isolation, matrix-bound nanovesicles and latex nanoparticles (Sigma Aldrich, L9904) were diluted at 1:10,000 and 1:100,000, respectively, in ultrapure water and quantified using a Malvern Panalytical Nanosight NS300 instrument (camera level = 11, syringe pump speed = 100, detection threshold = 5) to determine particle size and concentration.

### 2.5 Transmission electron microscopy

Matrix-bound nanovesicles diluted 1:10 in 1X PBS were gently pipetted onto the carbon-coated side of a 200 mesh square copper grid (Electron Microscopy Sciences, 50-260-38), and the sample was immediately wicked away using Whatman no. 1 filter paper. Following this, the grid was briefly stained with uranyl acetate twice and allowed to dry after removal of excess stain with filter paper. The grid was imaged with a JEOL 1230 transmission electron microscope and a Hamamatsu C4742-95 camera using AMT Image Capture version 7.0.0.268.

### 2.6 Isolation of whole cell lysate

At the end of the 21-day culture, fibroblast cell sheets were washed with 1X PBS and lysed in RIPA buffer (Thermo Fisher Scientific, 89900) supplemented with protease and phosphatase inhibitor (Thermo Fisher Scientific, A32961). Cell sheets in lysis buffer were then scraped from the culture dish, pipetted into a tube, and allowed to incubate on ice with intermittent vortexing. Whole cell lysates were aliquoted and stored at −80°C until further assays were performed.

### 2.7 Protein quantification and Western blotting

Total protein content of whole cell lysate and matrix-bound nanovesicles isolated from fibroblast cell sheets was quantified using a BCA assay (Thermo Fisher Scientific, 23227) according to manufacturer instructions. Matrix-bound nanovesicles were diluted 1:5 in 1X PBS, and whole cell lysate was diluted 1:20 in RIPA buffer. Bovine serum albumin standards were diluted in 1X PBS or RIPA buffer. Briefly, protein samples were combined with working reagent (50 parts Reagent A, 1 part Reagent B), incubated at 37°C for 30 minutes, and sample absorbance (562 nm) was analyzed using a plate reader (Infinite M Plex, TECAN). Sample protein concentration was calculated from a bovine serum albumin standard curve. Once quantified, whole cell lysate or matrix-bound nanovesicles were combined with 4X Laemmli buffer (BioRad, 1610747) supplemented with β-mercaptoethanol (Sigma Aldrich, M3148). Samples were then vortexed and boiled at 95°C for 5 minutes. Five micrograms of total protein were loaded in each well of a 10% Mini-PROTEAN TGX precast gel (BioRad, 4561034) along with a molecular weight reference ladder (Thermo Fisher Scientific, 26619). Gel electrophoresis was performed at 150 V for approximately 1 hour at room temperature in 1X running buffer (25 mM Tris, 190 mM Glycine, 0.1% SDS). Samples were transferred to a nitrocellulose membrane (BioRad, 1704158) at 2.5 A and up to 25 V for 3 minutes using a Trans-Blot Turbo Transfer System. After the transfer step, membranes were briefly washed in 1X PBS and blocked in 5% w/v non-fat dry milk dissolved in 1X PBS for one hour at room temperature with gentle shaking. Primary antibodies were diluted in 5% w/v bovine serum albumin in 1X PBS-T (0.2%), including anti-Integrin β1 (1:1,000, Cell Signaling Technology, 34971), anti-Alix (1:10,000, Proteintech, 12422-1-AP), and anti-CD63 (1:1,000, Proteintech, 25682-1-AP). Blots were incubated in the primary antibody overnight at 4°C with gentle shaking. The next day, membranes were washed in 1X PBS-T (0.2%) for five minutes three times to remove unbound antibody. Secondary antibody (Alexa Fluor 647 Goat anti-Rabbit IgG H+L, Invitrogen, A21244) was diluted in 5% w/v non-fat dry milk in 1X PBS-T (0.2%) at 1:10,000. Membranes were incubated in the dark for 1 hour at room temperature with gentle shaking. Following this, PBS-T washing steps were repeated and residual Tween-20 was removed with additional washes in 1X PBS. Blots were imaged using a BioRad ChemiDoc MP.

### 2.8 Fabrication of alginate hydrogels

Separate batches of Pronova UP VLVG (Sigma Aldrich, 42000501) (low-molecular weight) and UP MVG (Sigma, 42000101) (high-molecular weight) alginate were first purified via dialysis (3500 Da MWCO tubing) against Milli-Q water for four days. Alginate was then frozen, lyophilized, then dissolved in sterile FluoroBrite™ DMEM (Gibco, A1896701). Next, alginate solution was combined with MBVs or PBS in 1X PBS in a Luer lock syringe. A calcium sulfate slurry was mixed with 1X PBS and added to a second Luer lock syringe. These two syringes were then coupled, and the solutions were mixed several times. The hydrogel mixture was quickly deposited between two silanized glass plates to allow for gelation at room temperature (3-4 hours). An 8 mm biopsy punch (Thermo Fisher Scientific, 12460413) was used to cut several smaller hydrogel discs (∼100 µL), which were then transferred into a 48-well plate and incubated in 300 µL FluoroBrite™ DMEM at 37°C and 5% CO_2_. Final alginate and MBV concentrations were 2% w/v and 1 × 10^11^ particles/mL, respectively. Final calcium concentrations were 30, 40, and 60 mM for fast-relaxing hydrogels of 5, 10, and 20 kPa stiffnesses, respectively, and 10, 20, and 40 mM for slow-relaxing 5, 10, and 20 kPa gels, respectively.

### 2.9 Fluorescence labeling and imaging of matrix-bound nanovesicles in alginate hydrogels

Approximately 40 µL of MBV stock sample was combined with carboxyfluorescein succinimidyl ester (CFSE, 82.5 µM final concentration, Fisher Scientific, C1157) and incubated at 37°C for 20 minutes. Unbound CFSE was removed from the MBV suspension via ultrafiltration using a 30 kDa MWCO centrifugal filter (Sigma Aldrich, UFC503096). Samples in ultrafiltration columns were centrifuged at 14,000 g (4°C), the flow-through was discarded, and additional PBS was added to the retentate containing MBVs. This process was repeated until the retentate was clear. Fluorescent MBVs were retrieved from columns according to manufacturer instructions and encapsulated in alginate hydrogels for imaging as previously described (see Section 2.8). Hydrogels were imaged using a Zeiss Axio Observer 7 with ZEN 3.11 and an Axiocam 820.

### 2.10 Scanning electron microscopy of matrix-bound nanovesicles in alginate hydrogels

Alginate hydrogels with and without matrix-bound nanovesicles were fixed in 4% paraformaldehyde for 45 minutes at 37°C. After thorough washes with ultrapure water, fixed hydrogels were then dehydrated in an increasing series of ethanol concentrations (30%, 50%, 70%, 85%, 95%, 100%). The hydrogels were incubated in each ethanol concentration for 5 minutes, and each incubation was repeated three times. Following this, hydrogels were washed in a 1:1 solution of hexamethyldisilazane (HMDS, Sigma Aldrich, 379212) and 100% ethanol twice, then briefly rinsed with 100% HMDS. Lastly, hydrogels were placed in fresh HMDS and allowed to dry overnight in a chemical safety cabinet. Dehydrated hydrogels were sputter-coated with an SuPro Instruments ISC 150T Ion Sputter Coater and imaged with a Thermo Fisher Apreo C Scanning Electron Microscope.

### 2.11 Rheological properties of alginate hydrogels

Mechanical properties were characterized using an Anton Paar MCR 502 stress-controlled rheometer. The hydrogel sample was punched out and subsequently placed on the bottom plate, and an 8 mm parallel plate geometry was slowly lowered to make contact until a normal force 0.01 N was reached. First, a time sweep was performed at 1 Hz frequency and 1% strain to determine the storage (G’) and loss (G’’) moduli. From these data, the complex modulus (G*, Eq. 1) and the elastic modulus (Eq. 2) were calculated, assuming a Poisson’s ratio (ν) of 0.5. Upon reaching equilibrium in G’, a stress relaxation test was immediately initiated using a 10% step strain. The characteristic stress relaxation time was defined as the duration required for the stress to decay to half its maximum value.

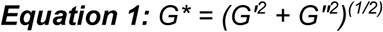

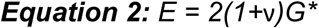

### 2.12 Quantification of matrix-bound nanovesicle release and retention

Alginate hydrogel discs (∼100 μL) with and without matrix-bound nanovesicles were incubated in 300 μL FluoroBrite DMEM in a 48-well plate and maintained at 37°C and 5% CO_2_. The culture medium surrounding alginate hydrogels was collected after 1, 4, 7, and 14 days of incubation and replaced with fresh medium. Prior to nanoparticle tracking analysis, the final volume of collected medium samples was adjusted to 1 mL with 50 mM EDTA to remove non-EV hydrogel particles that may have been released into the medium alongside MBVs. On day 14, hydrogel discs were chelated with 50 mM EDTA at room temperature with constant shaking, and the final volume of each hydrogel solution was further adjusted to 1 mL using 50 mM EDTA. Medium and chelated hydrogel samples were analyzed using a Malvern Panalytical Nanosight NS300 (camera level = 11, syringe pump speed = 100, detection threshold = 5) to measure released and retained MBVs, respectively. The percentage of retained (Eq. 3) and released (Eq. 4) particles were calculated using the following equations, with “X” representing a specific day of interest:

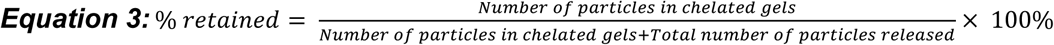

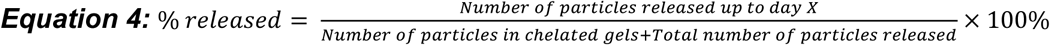

### 2.13 Quantification of latex nanoparticle release and retention

Amine-conjugated fluorescent latex nanoparticles (Sigma Aldrich, L9904) were encapsulated in alginate hydrogels as previously described (see Section 2.8). After 1, 4, 7, and 14 days of hydrogel incubation, the surrounding culture medium was collected, vortexed, and transferred to a black 96-well plate (Greiner Bio-One, 655209). Similarly, on day 14, hydrogels were chelated with 50 mM EDTA and the resulting solutions were transferred to a black plate. Medium and chelated hydrogel samples were analyzed using a plate reader (Infinite M Plex, TECAN) at excitation 475 nm and emission 540 nm with four reads per well. For release and retention experiments, instrument gain was set to 226 and 146, respectively.

### 2.14 Statistical Analysis

All data were analyzed in GraphPad Prism version 9.3.1 software using statistical tests described in the respective figure captions. Assumptions of normality and equal variance were assessed using Shapiro-Wilk and Brown-Forsythe tests, respectively, prior to statistical analyses. *P* values less than 0.05 were considered statistically significant for all data. Error bars are represented as mean ± standard deviation.

## 3. RESULTS

### 3.1 Human dermal fibroblasts secrete matrix-bound nanovesicles into extracellular matrix in vitro

MBVs extracted and purified from dermal fibroblast ECM were characterized for their quantity, size distribution, morphology, and surface marker expression (**Fig. 1A**). Nanoparticle tracking analysis confirmed that the isolated MBVs lay within the expected size distribution for small EVs (between 50-200 nm) with an average particle size of 139.9 nm and a concentration of 7.86 × 10^12^ particles/mL (**Fig. 1B**). Dermal fibroblast MBVs were additionally imaged using transmission electron microscopy to assess their morphology. Particles observed using TEM have deflated, EV-like morphologies (**Fig. 1C**). Lastly, we assessed MBV protein expression of putative EV biomarkers via Western blot analysis alongside whole cell lysate collected from dermal fibroblast cell sheets. We found that fibroblast MBVs expressed Alix, CD63, and Integrin β1 (**Fig. 1D, Supp. Fig. 1**). These targets have been suggested as positive EV biomarkers by International Society for Extracellular Vesicles’ most recent guidelines for minimal information for EV research ^35^.

**Figure 1:**
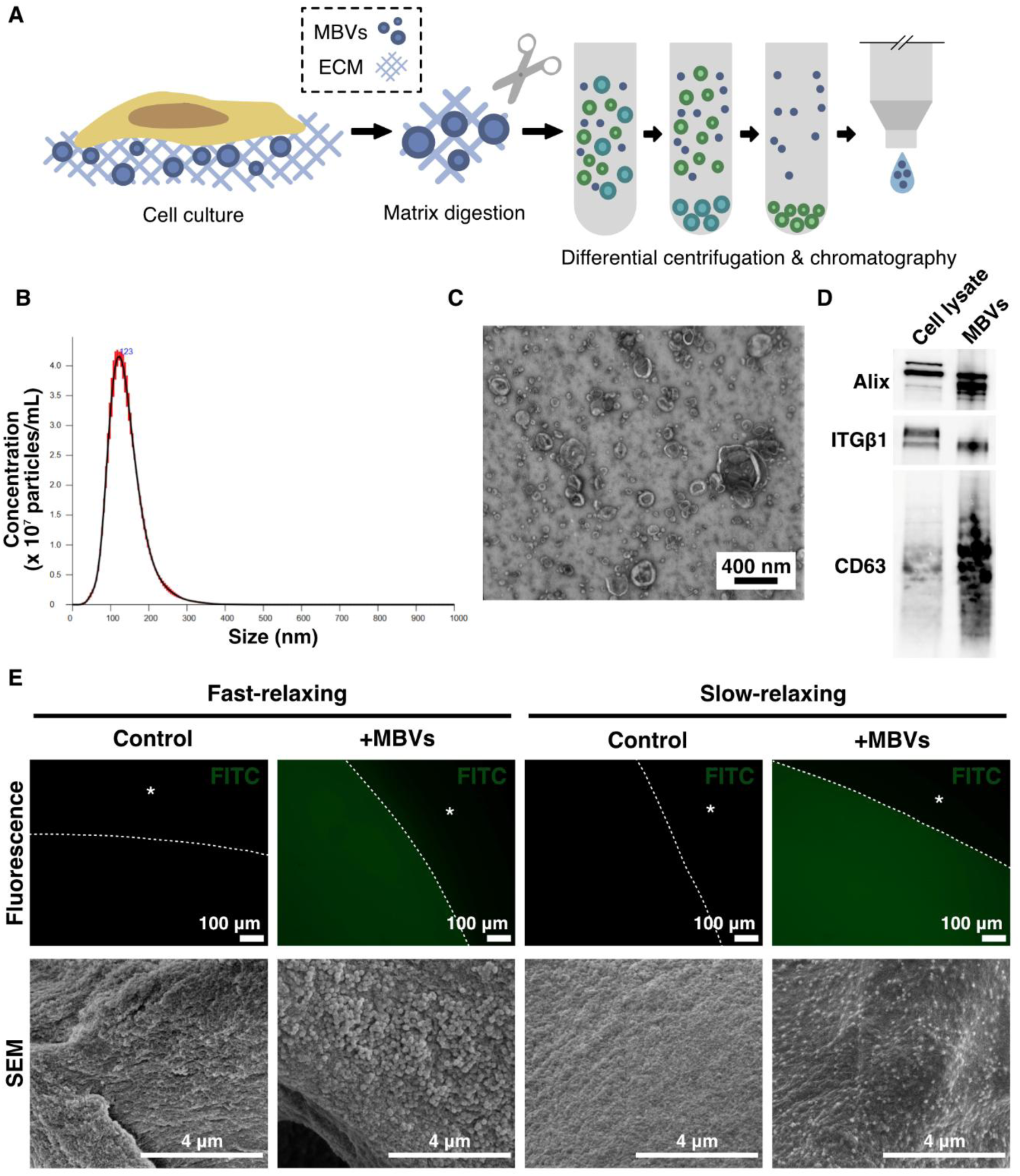
Isolation and characterization of MBVs from dermal fibroblasts and encapsulation in alginate hydrogels. **(A)** Schematic depiction of MBV isolation procedure from *in vitro* dermal fibroblast cultures. **(B)** Nanoparticle tracking analysis of diluted MBVs (1:10,000) measures particle size distribution and concentration. **(C)** Transmission electron micrograph of MBVs reveals vesicle-like morphology. **(D)** Western blot analysis of Alix (∼100kDa), Integrin β1 (∼100-130 kDa), and CD63 (∼35-55 kDa) expression in MBVs and dermal fibroblast cell lysate. **(E)** Fluorescence and scanning electron microscopy images of fast-relaxing and slow-relaxing alginate hydrogels with and without MBVs. *Top:* 5kPa hydrogels encapsulating CFSE-labeled MBVs, * denotes region outside of hydrogel. *Bottom:* 20kPa hydrogels encapsulating unlabeled MBVs.

We encapsulated MBVs in alginate hydrogels prior to crosslinking to ensure homogeneous MBV incorporation in hydrogels. Each hydrogel disc (∼100 µL) contained an estimated 1×10^10^ MBVs. To assess the distribution of MBVs in hydrogels using this fabrication method, MBVs fluorescently labeled with CFSE were encapsulated in alginate hydrogels and imaged. Fluorescence imaging demonstrated that MBVs were present throughout both slow- and fast-relaxing hydrogels, as there was a strong, uniform fluorescence signal that was exclusively within the bounds of the hydrogel. In contrast, hydrogels cast with vehicle control (1X PBS only) were non-fluorescent (**Fig. 1E**). Interactions between MBVs and alginate hydrogels were further visualized at higher magnification with scanning electron microscopy, and spherical, vesicle-like structures were observed in hydrogels (**Fig. 1E**).

### 3.2 Alginate hydrogels allow for independent control over stiffness and stress relaxation and MBVs do not impact matrix mechanical properties

Alginate-based 3D *in vitro* systems enable independent tuning of stiffness and stress relaxation without significantly altering matrix pore size or architecture^30^. Matrix stiffness in the alginate-based system was independently tuned by varying the concentration of the Ca^2+^ crosslinker (**Fig. 2A**)^29^. We optimized calcium concentrations to fabricate soft (5.61 ± 0.35 kPa), medium (10.07 ± 0.75 kPa), and stiff (20.38 ± 2.60 kPa) hydrogels (**Fig. 2B**). Stress relaxation, a measure of a material’s viscoelasticity, was tuned by varying the molecular weight of the alginate (**Fig. 2A**)^30^. We used low-molecular and high-molecular weight alginate to create fast- and slow-relaxing alginate hydrogels, respectively. For fast-relaxing alginate hydrogels, the half-time of stress relaxation (***τ***_1/2_) ranged from 70-180 seconds across the 5, 10, and 20 kPa stiffness conditions with a combined average of 160.2 ± 78.5 seconds. The ***τ***_1/2_ of slow-relaxing hydrogels ranged from 740 to 4,000 seconds across the same stiffness conditions with a combined average of 1,537 ± 776.6 seconds (**Fig. 2C, F**). We additionally performed stiffness and stress relaxation measurements on slow-relaxing, stiff gels with and without MBVs to assess whether MBVs affect the bulk mechanical properties of the alginate matrix (**Fig. 2D, E**). We found that there was no significant difference in either the stiffness (**Fig. 2D**) or stress relaxation (**Fig. 2E**) properties when MBVs were added to the alginate hydrogels.

**Figure 2:**
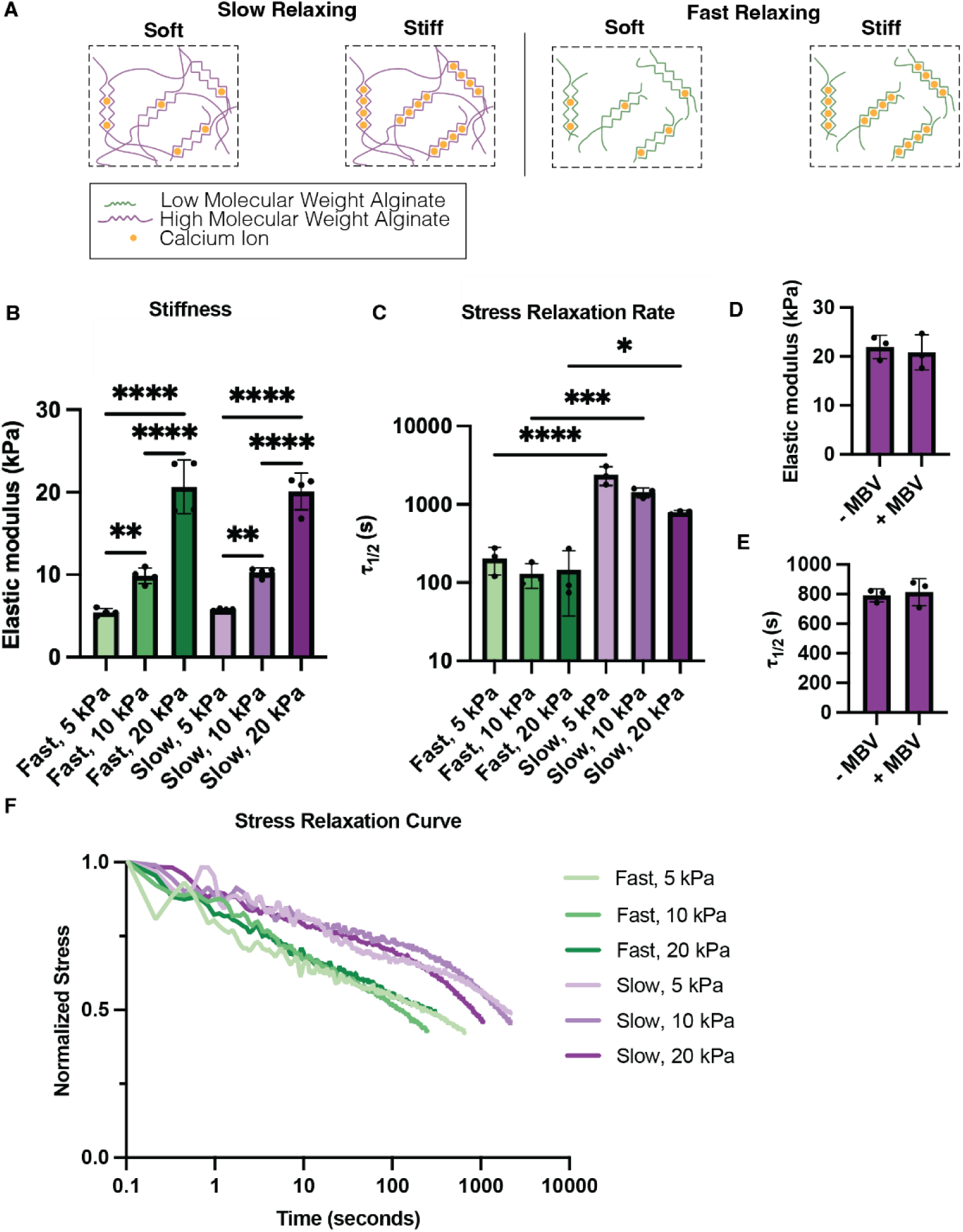
Independent tuning of alginate hydrogel stiffness and stress relaxation rate. **(A)** *Top*: Slow relaxing, or high molecular weight, alginate networks; stiffness can be tuned via calcium ion concentration. *Bottom:* Fast relaxing, or low molecular weight alginate networks; stiffness can be tuned via calcium ion concentration. **(B)** Elastic moduli of both fast and slow relaxing alginate hydrogels at stiffnesses 5, 10, and 20 kPa. **(C)** Stress relaxation rate of both fast and slow relaxing alginate hydrogels stiffnesses 5, 10, and 20 kPa. **(D)** Elastic moduli of slow relaxing, 20 kPa alginate hydrogels with (+) and without (−) MBV. **(E)** Stress relaxation rate of slow relaxing, 20 kPa alginate hydrogels with (+) and without (−) MBV. **(F)** Stress relaxation curve for both fast and slow relaxing alginate hydrogels at 5, 10, and 20 kPa. Significance was calculated using an unpaired t-test (D, E). If no statistical significance indicator bars are shown, there were no significant differences (i.e. p>0.05). *p<0.05, **p<0.01, ***p<0.001, and ****p<0.0001.

### 3.3 Stiff, fast-relaxing hydrogels exhibit increased MBV release compared to their softer counterparts

After verifying that bulk mechanical properties (stiffness and stress relaxation) of an alginate-based matrix could be independently tuned, and that MBVs do not significantly alter these properties, we next measured the release kinetics of MBVs from fast-relaxing alginate-based matrices (**Fig. 3A**). Prior to nanoparticle tracking analysis of released MBVs, we added EDTA to the collected culture medium to chelate non-EV nanoparticles that may co-release with MBVs over time, in order to minimize confounding variables in particle counts. Additionally, control hydrogels without MBVs were included in release and retention experiments to account for non-EV particles that were not removed by chelation. Analysis of release after 1 day demonstrated that MBV-loaded gels released significantly more particles compared to the vehicle control groups across stiffnesses; particle release from control hydrogels remained at least one order of magnitude below MBV-loaded hydrogels (**Supp. Fig. 2A**). When comparing MBV-loaded groups, the fast-relaxing, 20 kPa gels showed a significantly higher percentage of MBVs released compared to the 5 and 10 kPa groups on day 1 (**Fig. 3B, C**). Stiff 20kPa hydrogels continue to release more MBVs than their softer counterparts over time (days 1, 4, 7, and 14, **Fig. 3C-G**). In accordance with the increased release observed from the stiffest matrix, the fast-relaxing 5 and 10 kPa gels retained significantly more MBVs by day 14 compared to the 20 kPa group, with average MBV retention of 75.6 ± 3.4% (mean ± standard deviation) for 5 kPa gels, 77.1 ± 2.8% for 10 kPa gels, and 57.2 ± 2.5% for 20 kPa gels, and a combined average of 70.0 ± 9.8% retention for all stiffness groups (**Fig. 3H**). We did not find significant differences between non-EV particle numbers in the chelated control fast-relaxing gels, which were not loaded with any EVs, across all stiffnesses (**Supp. Fig. 2B**). Further, no significant differences in non-EV particle release were observed at each timepoint among the vehicle control fast-relaxing gels (5, 10, and 20 kPa) (**Supp. Fig. 2C**).

**Figure 3:**
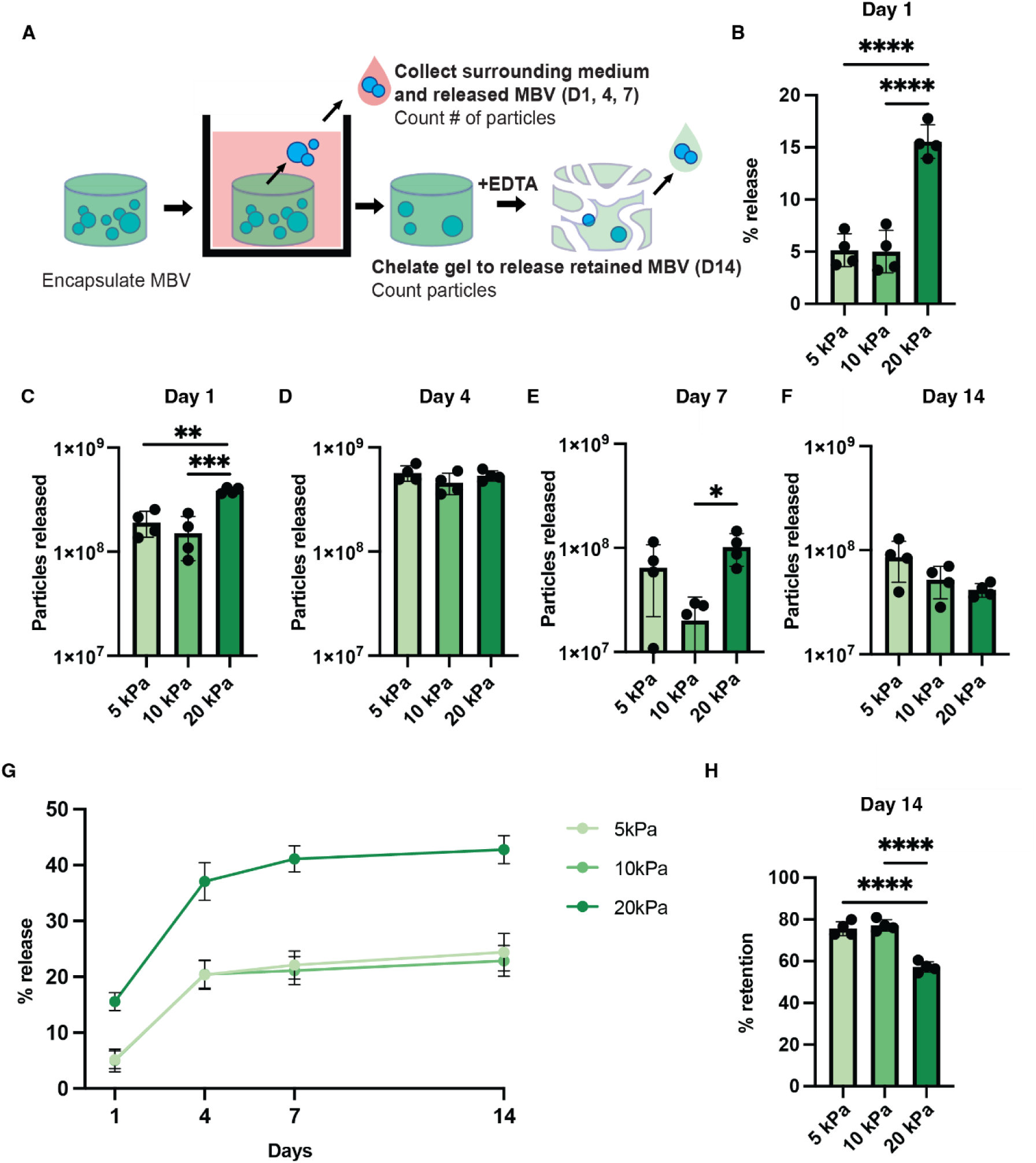
Stiffness-dependent MBV release from fast-relaxing alginate hydrogels. (A) Experimental procedure: MBV were encapsulated in fast-relaxing alginate hydrogels. Surrounding media was collected on days 1, 4, 7, and 14 to examine MBVs released and alginate hydrogels were chelated with EDTA on day 14 to examine MBVs retained. (B) Percent (%) of total MBV encapsulated released on day 1 by fast-relaxing alginate hydrogels at stiffnesses 5, 10, and 20 kPa. (C-F) Total MBV particles released by fast-relaxing alginate hydrogels at stiffnesses 5, 10 and 20 kPa on days 1, 4, 7, and 14. (G) Cumulative percent (%) release curve over the 14-day incubation period from fast-relaxing alginate hydrogels at stiffnesses 5, 10 and 20 kPa. (H) Percent (%) of total MBV encapsulated that were retained in the hydrogel on day 14 in fast-relaxing alginate hydrogels at stiffnesses 5, 10 and 20 kPa. Significance was determined using a one-way ANOVA followed by post-hoc multiple comparison tests for % release, particles released, and % retention. If no statistical significance indicator bars are shown, there were no significant differences (i.e. p>0.05). *p<0.05, **p<0.01, ***p<0.001, and ****p<0.0001.

### 3.4 Slow-relaxing hydrogels promote MBV retention in alginate hydrogels across several stiffnesses

We next characterized the release kinetics of MBVs from slow-relaxing alginate hydrogels (**Fig. 4A**). On day 1, MBV-loaded gels did not release significantly more particles than vehicle controls across all stiffnesses (5, 10, 20 kPa), indicating minimal MBV release from slow-relaxing gels (**Supp. Fig. 3A**). Furthermore, the percentage of total MBVs released on day 1 was not statistically different among the slow-relaxing 5, 10, and 20 kPa groups (**Fig. 4B**). Our analysis of MBV release from slow-relaxing gels over time revealed that the only significant difference in release between stiffness conditions occurred on day 4 between 5 and 10 kPa gels (**Fig. 4C-G**). On day 14, slow-relaxing gels across all stiffnesses demonstrated high MBV retention, with an average (± standard deviation) MBV retention of 96.3 ± 5.8% for 5 kPa gels, 97.0 ± 4.5% for 10 kPa gels, and 99.2 ± 0.7% for 20 kPa gels, and a combined average of 97.5 ± 4.0% retention (**Fig. 4H**). Critically, there was no significant difference in MBV retention based on gel stiffness (**Fig. 4H**). Additionally, we found no significant differences between non-EV particles measured in chelated control slow-relaxing hydrogels of varying stiffness (**Supp. Fig. 3B**). We also found no significant difference in release at each timepoint between control slow-relaxing 5, 10 and 20 kPa gels (**Supp. Fig. 3C**).

**Figure 4:**
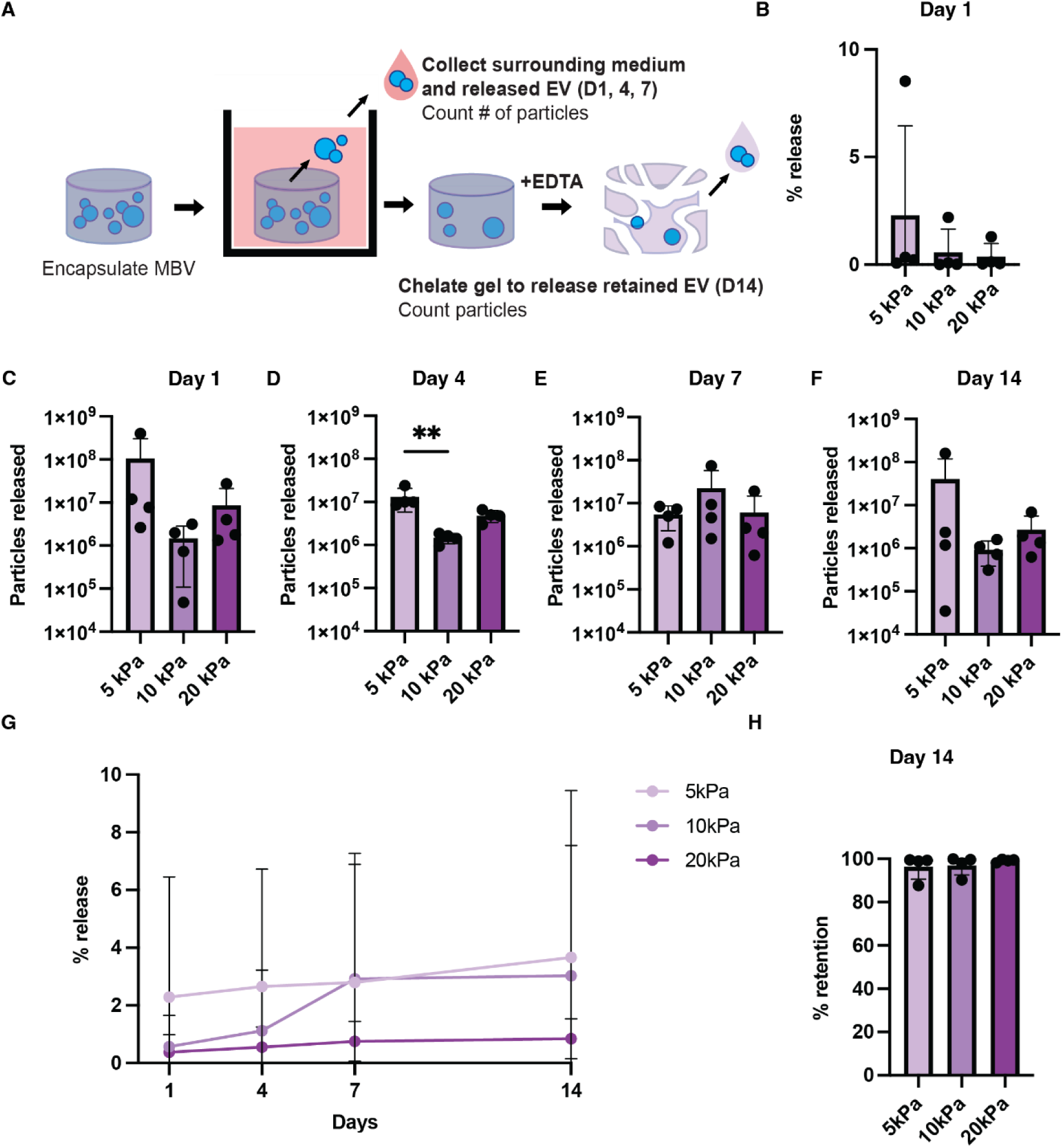
Stiffness-dependent MBV release from slow-relaxing alginate hydrogels. (A) Experimental procedure: MBV were encapsulated in slow-relaxing alginate hydrogels. Surrounding media was collected on days 1, 4, 7, and 14 to examine MBVs released and alginate hydrogels were chelated with EDTA on day 14 to examine MBVs retained. (B) Percent (%) of total MBV encapsulated released on day 1 by slow-relaxing alginate hydrogels at stiffnesses 5, 10, and 20 kPa. (C-F) Total MBV particles released by slow-relaxing alginate hydrogels at stiffnesses 5, 10 and 20 kPa on days 1, 4, 7, and 14. (G) Cumulative percent (%) release curve over the 14-day incubation period from slow-relaxing alginate hydrogels at stiffnesses 5, 10 and 20 kPa. (H) Percent (%) of total MBV encapsulated that were retained in the hydrogel on day 14 in slow-relaxing alginate hydrogels at stiffnesses 5, 10 and 20 kPa. Significance was determined using a Kruskal Wallis with Dunn’s multiple comparisons tests for % release, particles released, and % retention. If no statistical significance indicator bars are shown, there were no significant differences (i.e. p>0.05). *p<0.05.

### 3.5 Fast-relaxing hydrogels release more MBVs than slow-relaxing hydrogels

Analysis of MBV release kinetics revealed clear differences in initial release profiles between fast-and slow-relaxing alginate hydrogels (**Fig. 5A**). Specifically, fast-relaxing 20kPa hydrogels released significantly more MBVs on day 1 compared to their slow-relaxing counterparts at the same stiffness (**Fig. 5B**). Analysis of the cumulative MBV release profile from day 1 to 14 confirmed this trend, with fast-relaxing 5, 10, and 20 kPa hydrogels releasing more MBVs than their slow-relaxing counterparts over the 14-day period (**Fig. 5D-F**). Moreover, slow-relaxing 20 kPa gels retained a significantly higher percentage of MBVs by day 14 compared to their fast-relaxing equivalents (**Fig. 5C**).

**Figure 5:**
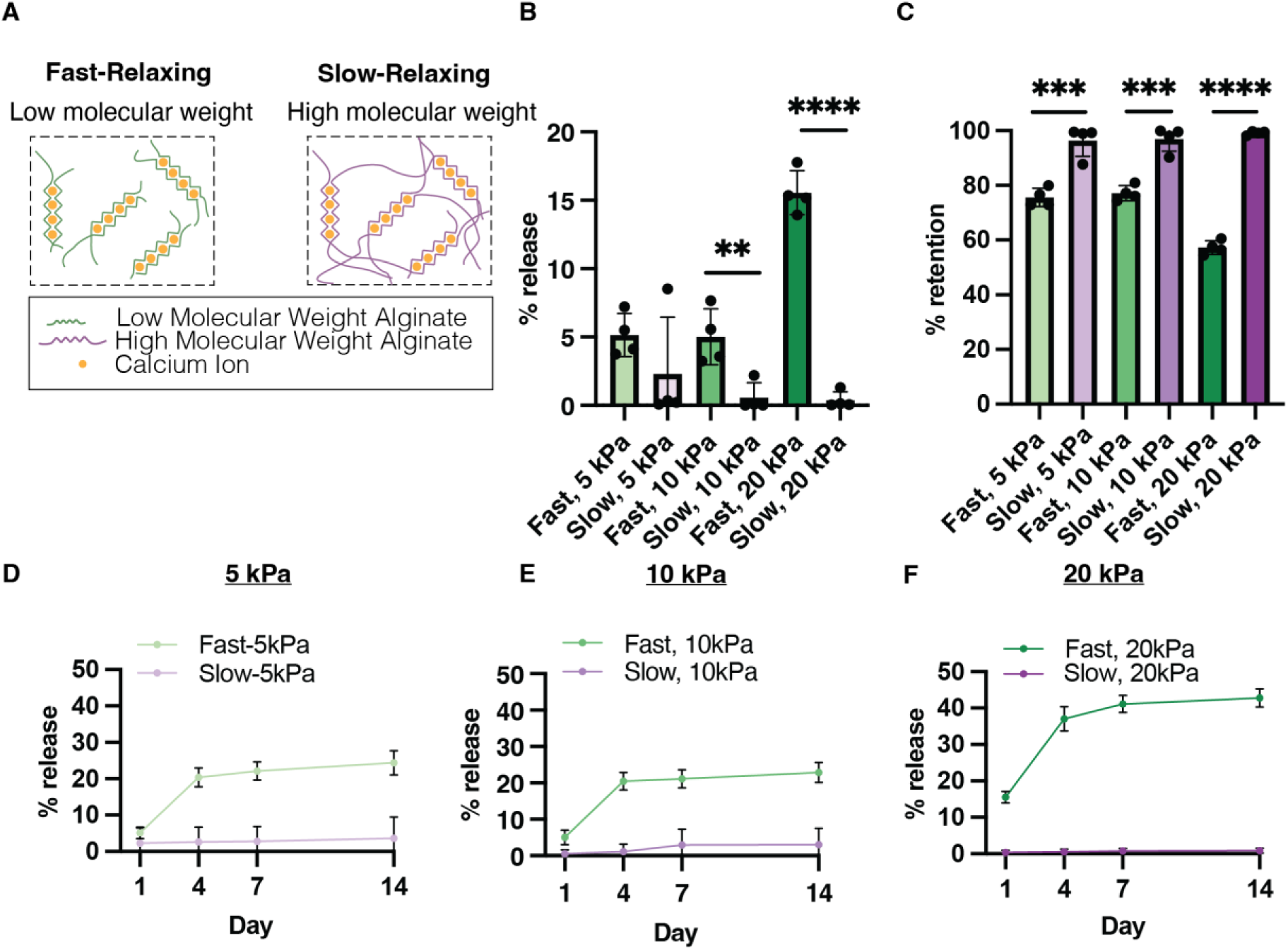
Fast-relaxing alginate hydrogels release more MBV than their slow-relaxing counterparts. (A) *Left:* Fast-relaxing, or low molecular weight, alginate hydrogel network. *Right:* Slow-relaxing, or high molecular weight, alginate hydrogel network. (B) Comparison of percent (%) of total MBV encapsulated released on day 1 by fast- and slow-relaxing alginate hydrogels at stiffnesses 5, 10 and 20 kPa. (C) Cumulative percent (%) release curve over the 14-day culture period from fast- and slow-relaxing alginate hydrogels at stiffness 5 kPa. (D) Cumulative percent (%) release curve over the 14-day culture period from fast- and slow-relaxing alginate hydrogels at stiffness 10 kPa. (E) Cumulative percent (%) release curve over the 14-day culture period from fast- and slow-relaxing alginate hydrogels at stiffness 20 kPa. (F) Comparison of percent (%) of total MBV encapsulated retained on day 14 by fast- and slow-relaxing alginate hydrogels at stiffnesses 5, 10 and 20 kPa. Significance was determined using an unpaired t-test or Mann-Whitney test for % release and % retention. If no statistical significance indicator bars are shown, there were no significant differences (i.e. p>0.05). **p<0.01.

### 3.6 Fast-relaxing hydrogels possess a time-dependent release profile

There was no significant difference in MBVs released from slow-relaxing 5, 10, or 20 kPa gels throughout days 1-14 (**Supp. Fig. 4A-C**). In contrast, the fast-relaxing hydrogels displayed a time-dependent release profile that varied by stiffness. There was significantly greater release on day 4 compared to days 1, 7 and 14 in fast-relaxing 5 kPa and 10 kPa gels (**Supp. Fig. 4D, E**). In fast-relaxing 20 kPa hydrogels, release was significantly greater on day 1 compared with days 7 and 14, and greatest on day 4 compared with all other time points (**Supp. Fig. 4F**).

### 3.7 Latex nanoparticles exhibit minimal release in alginate hydrogels regardless of matrix properties

We assessed whether nanoparticles of similar size to MBVs would exhibit similar stiffness- and stress relaxation-dependent release profiles in alginate hydrogels to elucidate whether interactions between MBVs and alginate matrices depend primarily on particle size. To determine this, we encapsulated commercially available fluorescent latex nanoparticles in fast- and slow-relaxing alginate hydrogels of varying stiffnesses (5, 10, 20 kPa) using a similar encapsulation procedure as MBVs and measured their release and retention (**Supp. Fig 5, Supp. Fig. 6**(−)). We found that latex nanoparticles averaged 83.7 nm in diameter in a similar size range to MBVs (**Supp. Fig. 5A**). Intriguingly, the earliest release event among all experimental conditions was detected on day 4 in soft (5 kPa), slow-relaxing hydrogels (**Supp. Fig. 6C**(−)), and these hydrogels continued to release latex nanoparticles until day 14 (**Supp. Fig. 6D, E**(−)). Similarly, only soft (5 kPa), fast-relaxing matrices exhibited latex nanoparticle release, though this occurred much later (day 14) relative to their slow-relaxing counterparts (**Supp. Fig. 5F**). Particle release from stiffer (10, 20 kPa) hydrogels was not detected over the 14-day incubation period regardless of matrix stress relaxation (**Supp. Fig. 5, 6**). Importantly, though some latex nanoparticle release was detected in alginate hydrogels, there were no significant stiffness-dependent differences in particle retention in either fast- or slow-relaxing hydrogels, indicating minimal release throughout the 14-day incubation period regardless of matrix mechanical properties (**Supp. Fig. 5C, 6B**).

## 4. DISCUSSION

MBVs are promising EVs for regenerative medicine applications, but like other EV types, MBVs are most frequently administered as a bolus injection *in vivo* and suffer from poor persistence and organ localization. Local intramuscular and subcutaneous MBV injections have longer retention times at the original site of administration (3 and 7 days, respectively) compared to IV injections, but MBVs also localize to the kidneys within the first day of injection^10^. While this may be longer than retention times for other types of EVs (24 hours), this biodistribution study suggests that current routes of MBV administration are largely inefficient and untargeted. Most importantly, the premature systemic clearance of MBVs, as with all EVs, is suboptimal for the treatment of conditions requiring weeks-to-months of therapeutic activity, such as large-scale bone defect repair^15^. Alginate hydrogels have been widely used to sequester EVs in a wound site *in vivo* with improved treatment outcomes^36–39^, yet it remains poorly understood how these hydrogels can be engineered to achieve specific therapeutic timescales. To help address this limitation and inform EV delivery strategies, we explored how the mechanical properties of alginate hydrogels can be tuned to control MBV release.

Human dermal fibroblasts were cultured *in vitro* with L-ascorbic acid to increase cellular proliferation and collagen deposition^40,41^, thus promoting MBV accumulation in the cell-deposited extracellular matrix^5^. As with any tissue-derived EV, the extracellular matrix must first be dissociated to release MBVs prior to enrichment with conventional EV isolation techniques. The use of Liberase TH for enzymatic digestion and differential centrifugation and size-exclusion chromatography for enrichment of MBVs was motivated by previous studies reporting higher particle yields and bioactivity with such combination of methods^42^. We show that enriched dermal fibroblast MBVs express Alix, Integrin β1, and CD63, which fall under Categories 1 and 2 of EV protein markers suggested by MISEV 2023^35^. While others have shown 3T3 fibroblast MBVs to express CD63^5^, we are the first to show that Alix and Integrin β1 may be potential biomarkers for fibroblast MBVs. This aligns with previous data showing that mesenchymal stem cell MBVs likewise are enriched in Alix, Integrin β1, and CD63 compared to other EV types^7^. Taken together, the combination of Western blotting, electron microscopy, and nanoparticle tracking analysis strongly indicated the presence of EVs in MBV preparations. We then encapsulated MBVs into alginate hydrogels and further verified MBV presence and distribution using fluorescence microscopy and scanning electron microscopy. Importantly, matrix stiffness and stress relaxation were unaltered by the incorporation of MBVs into the hydrogels; these results are in agreement with previous characterization of exosome-loaded hydrogels compared to their non-exosome loaded counterparts^43^.

We show that fast-relaxing hydrogels are characterized by the release of MBVs within the first 4 days of incubation in culture medium, and hydrogel stiffness determines the extent of this release. These results are consistent with previous findings, in which relatively stiff alginate hydrogels (3 kPa) released more liquid-EVs in a 24-hour period compared to their soft counterparts (500 Pa) and this correlation was observed across liquid-EVs isolated from multiple cell types^16^. This trend could be explained by the dynamic nature of alginate egg-box junctions; while higher calcium concentrations increase crosslink density, the reversible and non-permanent nature of these ionic bonds allows for frequent dissociation and re-association. This high rate of exchange at saturated coordination sites may facilitate localized network rearrangements, ultimately promoting the diffusion of MBVs out of the hydrogels. Additionally, in physiological buffers such as DMEM, monovalent cations (e.g. Na^+^) compete with crosslinking divalent cations (e.g. Ca^2+^) in alginate hydrogels, leading to calcium dissociation^44,45^, which could coincide with MBV release. Regardless, the unique structure and crosslinking mechanism of these alginate hydrogels give rise to matrix stiffness-dependent MBV release profiles that may not be observed in material systems of different composition, structure, and crosslinking mechanism. For instance, others have reported that soft hydrogels release EVs more rapidly than stiff hydrogels in hyaluronic acid methacrylate scaffolds^18^, though the numbers of EVs were not measured.

In contrast with fast-relaxing hydrogels, slow-relaxing hydrogels exhibited exceptional long-term MBV sequestration regardless of stiffness in our studies, with a combined average of 97.5% particle retention after 14 days. These results are supported by other studies that have shown prolonged matrix stress relaxation impeded EV release^16,17^. Covalently crosslinked alginate hydrogels, which exhibit elastic behavior, have previously been shown to release more EVs than their stress relaxing, physically crosslinked counterparts in 24 hours^16^. In a different material system, a reinforcing polycaprolactone mesh was used to increase the stress relaxation time constant of GelMA hydrogels, and total EV release decreased from 60% to 12% over 14 days^17^. Though a similar relationship between matrix stress relaxation and EV release was observed in our experiments, the slow-relaxing alginate hydrogels utilized herein released a much lower percentage of EVs over a 14-day period across all stiffness conditions. In our experiments, a lack of MBV release from slow-relaxing hydrogels could be attributed to an increase in chain length and a decrease in chain mobility that obstructs MBV transport in hydrogels in contrast with fast-relaxing matrices that exhibit fewer chain entanglements and may more easily undergo network remodeling^30,46–48^. Importantly, given that no stiffness-dependent differences in MBV release were observed in slow-relaxing hydrogels unlike their fast-relaxing counterparts, we found that matrix stress relaxation is a significant driver of MBV sequestration and overcomes the effects of matrix stiffness on MBV release.

We also assessed whether interactions between MBVs and alginate matrices were solely dependent on particle diameter by using 100 nm latex nanoparticles as a size control in release experiments. Our intent in using latex particles was to establish a rigid, size-matched physical benchmark to compare to MBV release. By comparing the release kinetics of these rigid synthetic spheres to the more deformable MBVs, we aimed to isolate whether the transport was governed solely by particle size. Specifically, using alginate-calcium matrices allows us to alter mechanical properties without changing the pore size of the matrix^32^. We showed that observed trends in MBV release and retention in alginate matrices were not upheld with latex nanoparticles as they were primarily sequestered in hydrogels regardless of matrix mechanical properties. These results were unsurprising, as latex nanoparticles are relatively rigid and lack the deformability of MBVs that may enable escape from the confining alginate network. Others have shown a similar absence of particle diffusion with gold nanoparticles encapsulated in alginate interpenetrating networks with pore sizes less than the particle diameter^49^. Macromolecules such as bovine serum albumin and dextran can diffuse out of alginate matrices, but unlike MBVs, this diffusion is unaffected by changes in matrix stiffness^50^. Similarly, the release profile of liposomes of comparable size and lipid content to EVs from alginate hydrogels is unaffected by matrix stiffness^16^. The unique interactions between MBVs and alginate matrices could be attributed to the complex features of EVs, such as surface charge, membrane deformability, an aqueous core, and numerous surface proteins and lipids, that cannot be approximated with macromolecules and synthetic nanoparticles. Thus, the combination of MBVs and hydrogels is additionally advantageous in that material mechanical properties can be engineered to fine-tune the extent of MBV release or to promote their retention, unlike non-EV particles.

Our study supports the notion that hydrogels offer a solution to current challenges associated with EV administration as these materials can be specifically designed to achieve EV delivery profiles tailored to their intended clinical applications. Current hydrogel design strategies to modulate EV release primarily leverage controlled material crosslinking and degradation mechanisms or non-specific and specific molecular interactions between EVs and the material. Hydrogels can be designed to respond to stimuli such as pH^20^, temperature^51^, and light^22^ to release EVs, and EVs can also be immobilized to materials by covalent attachment^52,53^, electrostatic interactions^23,24^, or integrin-binding peptide sequences^54–56^. Unfortunately, many of these existing studies measure EV release only up to a week or less^21,57,58^, and many use indirect measurements of EV release, such as total protein concentration^51,57–61^, that fail to account for the inherent heterogeneity of EV populations. In certain cases, EV release is not measured at all^62^. Many of these materials require relatively complicated modification strategies to promote release or binding of EVs, such as the use of click chemistry^53,60^ and photocleavable linkers^22^. Most importantly, we found that few studies seek to pinpoint the effects of specific, isolated material properties on EV interactions with hydrogels, and in some cases, the material platforms used may possess several characteristics that affect their interactions with EVs. We addressed some of these limitations by utilizing unmodified alginate hydrogels that are relatively simple to fabricate with varying viscoelastic properties and measuring EV release and retention over a prolonged, 14-day time period. Additionally, while many studies have incorporated various types of EVs in engineered hydrogels toward diverse clinical applications, relatively few have investigated the combination of MBVs and hydrogels^6,63^ despite the evidenced anti-inflammatory properties of MBVs that may be useful in biomaterials-based tissue engineering strategies. Further, to date, none have interrogated interactions between MBVs and hydrogel matrices to control MBV delivery, making our study one of the first to embed MBVs in hydrogels and the first to decouple the effects of both matrix stiffness and stress relaxation on long-term MBV release and retention in alginate hydrogels.

Previous alginate hydrogel-based EV delivery systems have demonstrated substantial variability in EV release kinetics, with one study reporting near-complete release as early as 5 days^64^ and others delaying complete release until day 10^65,66^. Sustained release over 14 days, similar to that observed in our system, has also been described, although EV release and retention rates are much poorer, with one study showing 85% cumulative release^67^ and another achieving 38% retention^68^ in alginate hydrogels by day 14. By merely prolonging the stress relaxation of alginate matrices without further material modifications, we were able to achieve an average of 97.5% EV retention in slow-relaxing hydrogels over 14 days; such high EV retention has not been accomplished even with RGD-modified alginates engineered to promote EV tethering, which only retain up to 80% of EVs in one week^55^.

From a translational perspective, our findings suggest that fast-relaxing matrices may be more suitable towards therapeutic design strategies of biomaterials for early MBV release from the site of administration. In contrast, slow-relaxing matrices may better support the development of therapeutic designs of biomaterials for extended MBV sequestration that promote prolonged MBV tissue localization and therapeutic timelines. The therapeutic dose of EVs is equally critical to their bioavailability in clinical applications, and we further show that the extent of MBV release can be fine-tuned by modulating the stiffness of fast-relaxing matrices. However, it is well known that matrix mechanical properties inform cellular behavior through dynamic reciprocity, with stiffness and stress relaxation influencing fundamental processes such as cell phenotype, migration, epigenetic modifications or matrix deposition^30,69–76^. Consequently, the stiffness and stress relaxation of implanted MBV-laden hydrogels may also impact the surrounding tissue microenvironment beyond the spatiotemporal control of MBVs. Due to their relatively recent discovery, it remains unclear whether MBVs override the effect of matrix mechanical stimuli on cell response, much less at prolonged timescales. Nonetheless, a careful balance must be struck so that hydrogel platforms can simultaneously support cellular functions and the controlled release of MBVs.

## 5. CONCLUSION

Matrix-bound nanovesicles (MBVs) are a type of matrix-resident extracellular vesicle with unique anti-inflammatory properties in clinical applications; however, leveraging these properties for clinical use is often hindered by a lack of fundamental understanding regarding their controlled delivery. Here, we show that alginate hydrogels present an ideal MBV delivery vehicle as material stiffness and stress relaxation can be modulated to more precisely control MBV release and retention, and importantly, can be tuned to retain nearly 100% of MBVs over a two-week period. We are the first to encapsulate MBVs in alginate hydrogels and demonstrate that stiffer fast-relaxing hydrogels result in greater MBV release, and slow-relaxing hydrogels can largely retain MBVs regardless of stiffness changes. Our results highlight unique interactions between MBVs and hydrogel matrices and introduce material mechanical property modulation as a strategy for controlling MBV delivery in therapeutic applications.

## ACKNOWLEDGMENTS

This work was supported by the National Science Foundation Materials Research Science and Engineering Centers (Division of Materials Research, DMR2308708). This work was also partially supported by a National Institutes of Health T32 Training Grant (1T32GM141846) (R.D.M., J.A.B., G.M.G). The content is solely the responsibility of the authors and does not necessarily represent the official views of the National Science Foundation or the National Institutes of Health. This study utilized instrumentation provided by the University of California-Santa Barbara’s Biological Nanostructures Laboratory, the Neuroscience Research Institute-Department of Molecular, Cellular and Developmental Biology Microscopy Facility, and the Materials Microscopy and Microanalysis Facility. The authors would like to thank Drs. Carolyn Mills and Beth Pruitt for granting our use of a BioRad Chemidoc MP and Zeiss Axio Observer 7, respectively, for the completion of this study.

## DISCLOSURE OF INTEREST

The authors have no conflicts of interest to disclose.

## AUTHOR CONTRIBUTIONS

**Renata Dos Reis Marques:** Formal analysis, Investigation, Methodology, Visualization, Validation, Writing – Original Draft Preparation, Writing – Review & Editing. **Jane Baude:** Formal analysis, Investigation, Methodology, Visualization, Validation, Writing – Original Draft Preparation, Writing – Review & Editing. **Gianna Gathman:** Investigation. **Ava Salami:** Investigation. **Ryan Stowers:** Funding Acquisition, Project Administration, Resources, Supervision, Writing – Review & Editing. **Marley Dewey:** Funding Acquisition, Project Administration, Resources, Supervision, Writing – Review & Editing.

## Notes

### Competing Interest Statement

The authors have declared no competing interest.

