## Supplementary files for "Matrix stiffness and stress relaxation regulate matrix-bound nanovesicle release from alginate hydrogels"

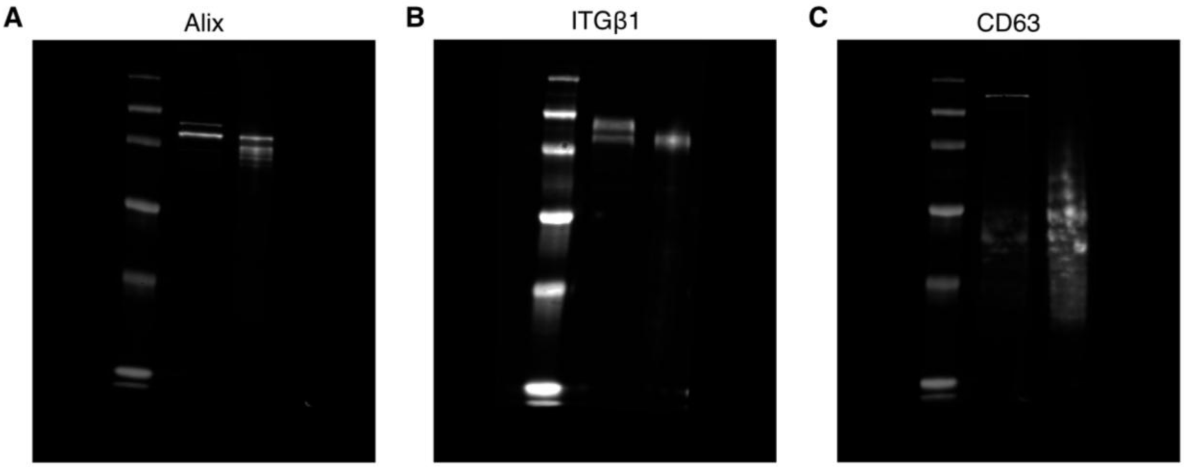


**Supplementary Figure 1:** Raw Western blot images of dermal fibroblast cells and MBVs. **(A)** Alix. **(B)** Integrin β1. **(C)** CD63.

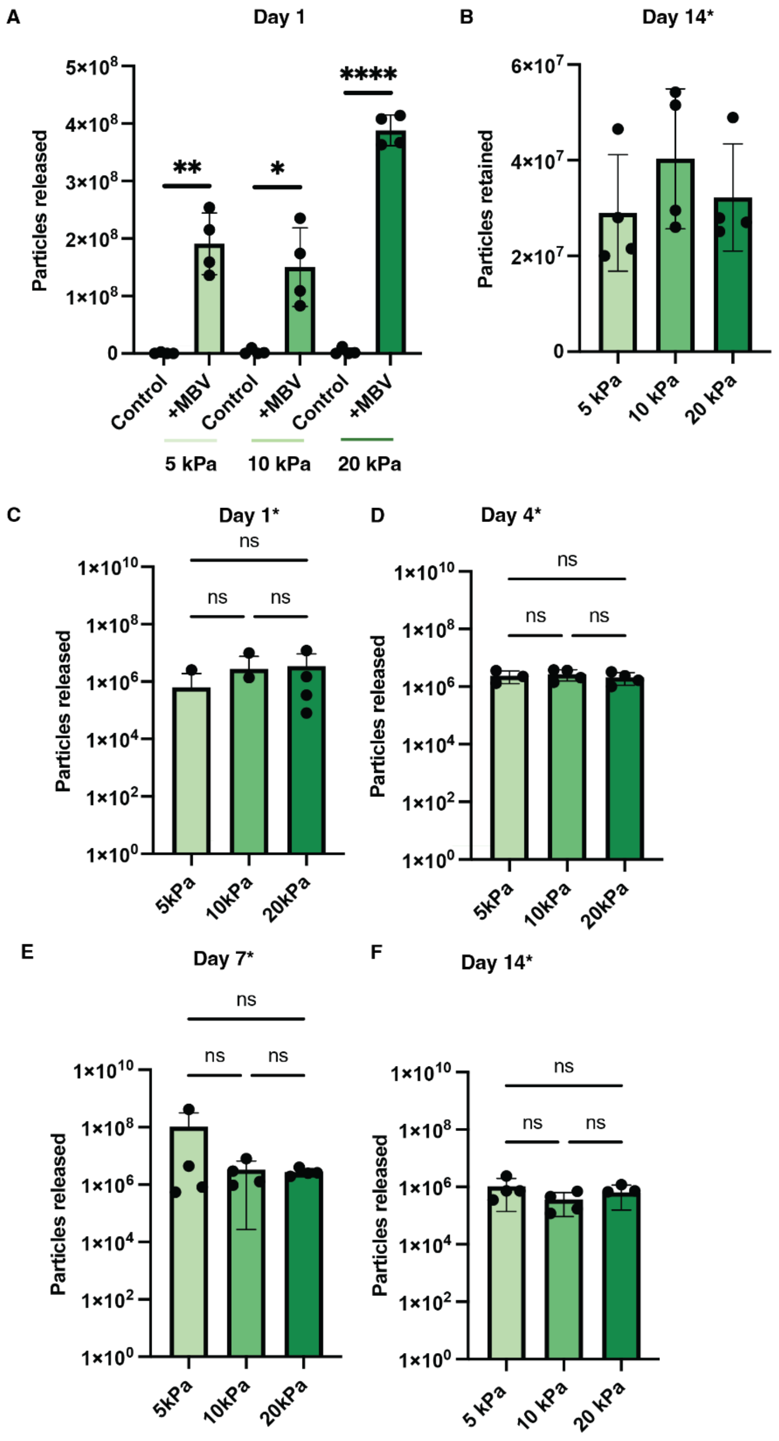


**Supplementary Figure 2: Particle release from control* fast-relaxing alginate hydrogels. (A)** Comparison of particles released on day 1 from PBS (control) versus MBV loaded fast-relaxing alginate hydrogels at stiffnesses 5, 10, and 20 kPa. **(B)** Non-EV particles retained on day 14 by fast-relaxing alginate hydrogels at stiffnesses 5, 10, and 20 kPa. **(C-F)** Non-EV particles released from fast-relaxing alginate hydrogels at stiffnesses 5, 10 and 20 kPa after **(C)** 1 day, **(D)** 4 days, **(E)** 7 days, and **(F)** 14 days of incubation. Significance was determined using a Welch's t-test (A), a one-way ANOVA followed by Tukey's post-hoc multiple comparisons test (B, D, F), and a Kruskal Wallis followed by a Dunn's multiple comparisons test (C, E). If no statistical significance indicator bars are shown, there were no significant differences ($p>0.05$). *$p<0.05$, **$p<0.01$, and ****$p<0.0001$.

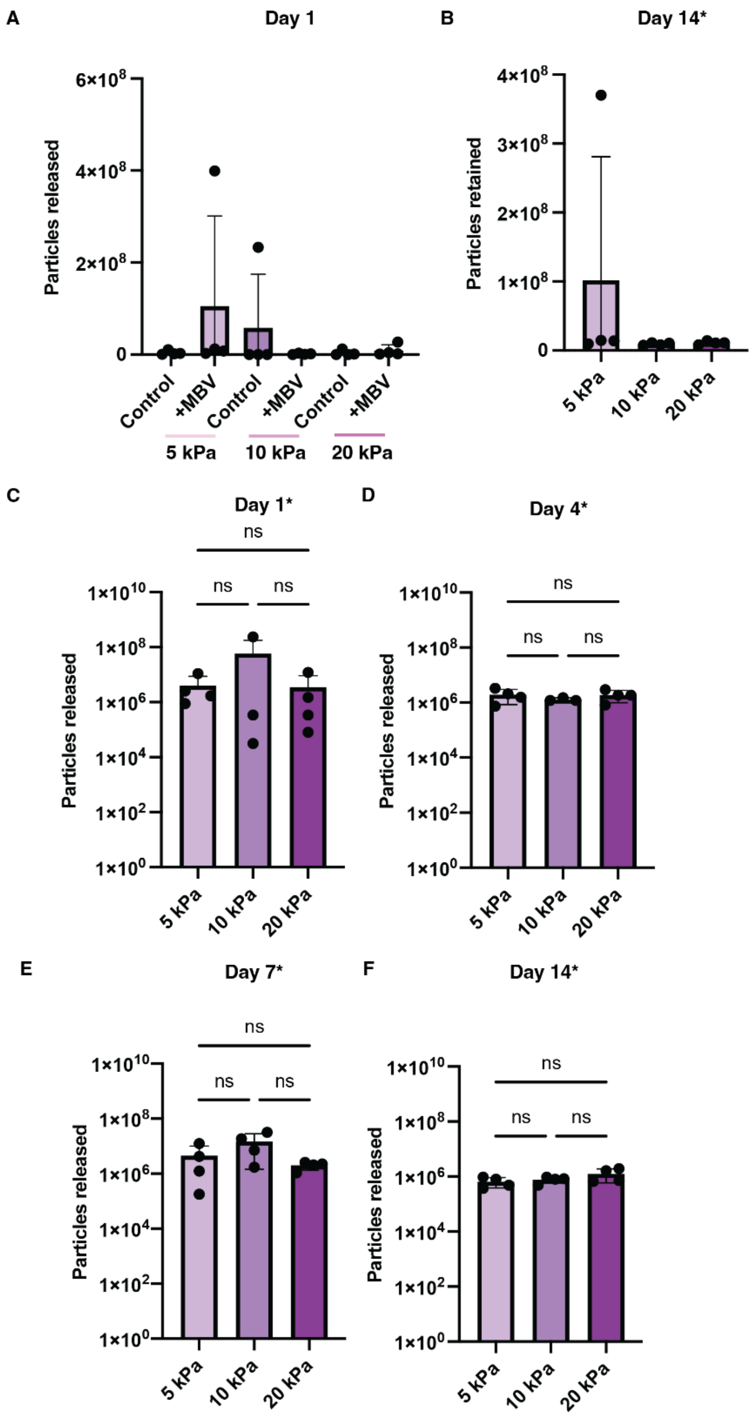


**Supplementary Figure 3: Particle release from PBS (control*) slow-relaxing alginate hydrogels. (A)** Comparison of particles released on day 1 from PBS (control) versus MBV loaded slow-relaxing alginate hydrogels at stiffnesses 5, 10, and 20 kPa. **(B)** Non-EV particles retained on day 14 by slow-relaxing alginate hydrogels at stiffnesses 5, 10, and 20 kPa. **(C-F)** Non-EV particles released from slow-relaxing alginate hydrogels at stiffnesses 5, 10 and 20 kPa after **(C)** 1 day, **(D)** 4 days, **(E)** 7 days, and **(F)** 14 days of incubation. Significance was determined using a Mann-Whitney test (A), a Kruskal-Wallis followed by a Dunn's multiple comparisons test (B, C), a one-way ANOVA followed by Tukey's post-hoc multiple comparisons test (D), and a Brown-Forsythe and Welch ANOVA followed by a Dunnett's T3 multiple comparisons test (E, F). If no statistical significance indicator bars are shown, there were no significant differences ($p>0.05$).

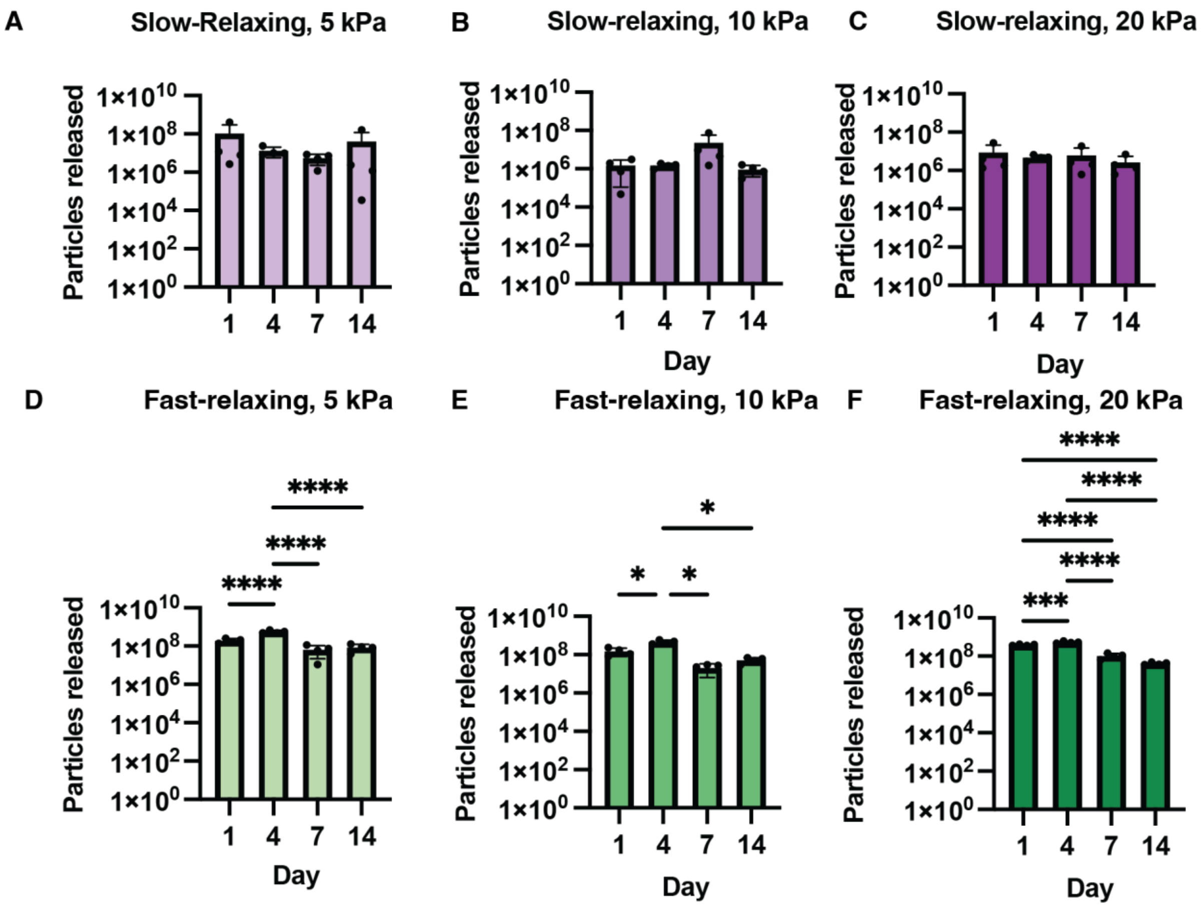


**Supplementary Figure 4: Comparison of EV release within each alginate hydrogel condition by day. (A-C)** Comparison of particles released on days 1, 4, 7, and 14 from slow-relaxing alginate hydrogels of **(A)** 5 kPa, **(B)** 10 kPa, and **(C)** 20 kPa stiffness. **(D-F)** Comparison of particles released on days 1, 4, 7, and 14 from fast-relaxing alginate hydrogels at **(D)** 5 kPa, **(E)** 10 kPa, and **(F)** 20 kPa stiffness. Significance was determined using a Kruskal Wallis followed by a Dunn's multiple comparisons test (A-C), a one-way ANOVA followed by Tukey's post-hoc multiple comparisons test (D, F), or a Brown-Forsythe and Welch ANOVA followed by a Dunnett's T3 multiple comparisons test (E). If no statistical significance indicator bars are shown, there were no significant differences ($p>0.05$). *$p<0.05$, ***$p<0.001$, and ****$p<0.0001$.

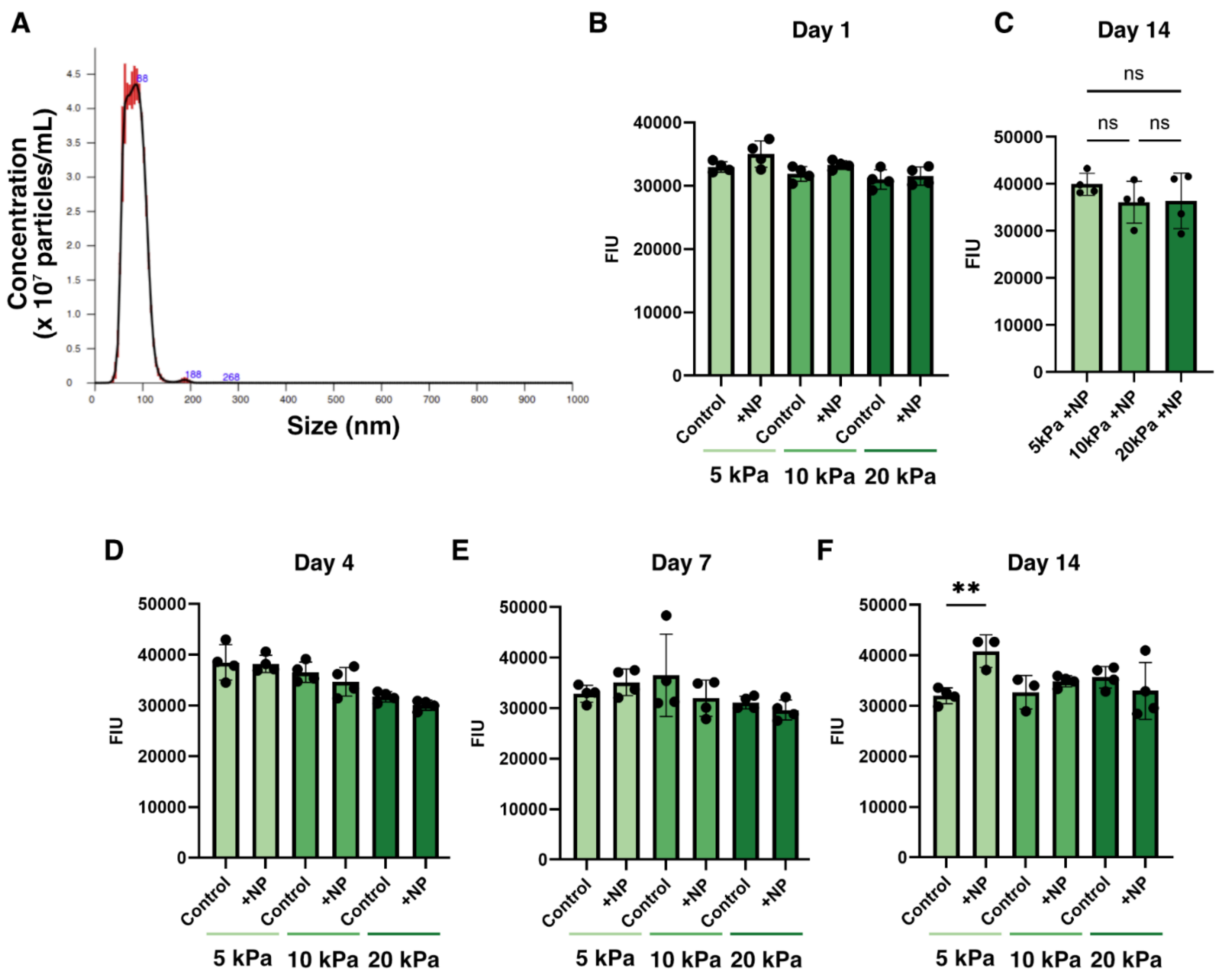


**Supplementary Figure 5: Latex nanoparticle (NP) release from fast-relaxing alginate hydrogels. (A)** Nanoparticle tracking analysis of latex NPs reveals a similar particle size to MBVs. **(B)** Comparison between fluorescence intensity units (FIU) of culture medium collected from control or NP-loaded fast-relaxing hydrogels of 5, 10, and 20 kPa stiffness after 1 day of incubation. **(C)** Comparison between FIU of chelated fast-relaxing hydrogels of 5, 10, and 20 kPa stiffness after 14 days of incubation (retention). **(D-F)** Comparison between FIU of culture medium collected from control or NP-loaded fast-relaxing hydrogels at 5, 10, and 20kPa stiffness after **(D)** 4 days, **(E)** 7 days, or **(F)** 14 days of incubation. Significance was determined using an unpaired t-test (B, D, E, F) or a one-way ANOVA followed by a Tukey's post-hoc multiple comparisons test (C). If no statistical significance indicator bars are shown, there were no significant differences ($p>0.05$). **$p<0.01$.

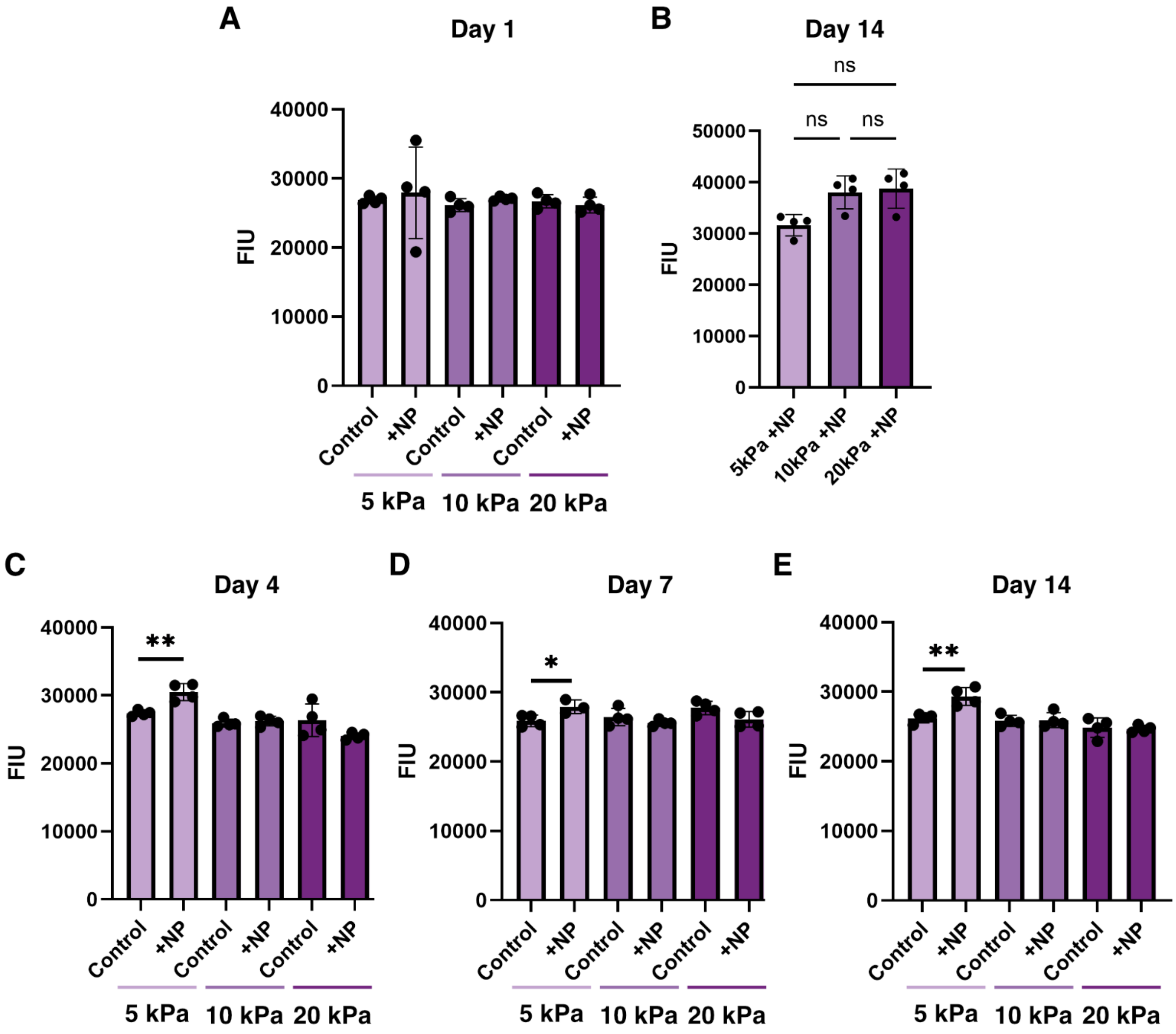


**Supplementary Figure 6: Latex nanoparticle (NP) release from slow-relaxing alginate hydrogels. (A)** Comparison between fluorescence intensity units (FIU) of culture medium collected from control or NP-loaded slow-relaxing hydrogels of 5, 10, and 20 kPa stiffness after 1 day of incubation. **(B)** Comparison between FIU of chelated slow-relaxing hydrogels of 5, 10, and 20 kPa stiffness after 14 days of incubation (retention). **(C-E)** Comparison between FIU of culture medium collected from control or NP-loaded slow-relaxing hydrogels of 5, 10, and 20kPa stiffness after **(C)** 4 days, **(D)** 7 days, or **(E)** 14 days of incubation. Significance was determined using a Mann-Whitney test (A-5 kPa), unpaired t-test (A-10 and 20 kPa, C-5 and 10 kPa, D, E), Welch's t-test (D- 20 kPa), or a Kruskal-Wallis followed by a Dunn's multiple comparisons test (B). If no statistical significance indicator bars are shown, there were no significant differences ($p>0.05$). *$p<0.05$ and **$p<0.01$.
